# Dual-view light-sheet imaging through tilted glass interface using a deformable mirror

**DOI:** 10.1101/2020.10.20.345306

**Authors:** N Vladimirov, F Preusser, J Wisniewski, Z Yaniv, RA Desai, A Woehler, S Preibisch

## Abstract

Light-sheet microscopy has become one of the primary tools for imaging live developing organisms because of its high speed, low phototoxicity, and optical sectioning capabilities. Detection from multiple sides (multi-view imaging) additionally allows nearly isotropic resolution via computational merging of the views. However, conventional light-sheet microscopes require that the sample is suspended in a gel to allow optical access from two or more sides. At the same time, the use of microfluidic devices is highly desirable for many experiments, but geometric constrains and strong optical aberrations caused by the coverslip titled relative to objectives make the use of multi-view lightsheet challenging for microfluidics.

In this paper we describe the use of adaptive optics (AO) to enable multi-view light-sheet microscopy in such microfluidic setup by correcting optical aberrations introduced by the tilted coverslip. The optimal shape of deformable mirror is computed by an iterative stochastic gradient-descent algorithm that optimizes PSF in two orthogonal planes simultaneously. Simultaneous AO correction in two optical arms is achieved via a knife-edge mirror that splits excitation path and combines the detection path.

We characterize the performance of this novel microscope setup and, by dual-view light-sheet imaging of *C.elegans* inside a microfluidic channel, demonstrate a drastic improvement of image quality due to AO and dual-view reconstruction. Our microscope design allows multi-view light-sheet microscopy with microfluidic devices for precisely controlled experimental conditions and high-content screening.

## Introduction

Over the last decade, light-sheet fluorescence microscopy (LSFM, also known as selective-plane illumination microscopy, SPIM) has become one of the primary tools for imaging live developing organisms due to its low photo-toxicity, optical sectioning, isotropic resolution and high speed (Huisken, Swoger, Del Bene, Wittbrodt, & Stelzer, 2004; Keller, Schmidt, Wittbrodt, & Stelzer, 2008). A conventional light-sheet microscope uses two objectives orthogonal to each other: one creates light-sheet excitation, while the other detects light emitted by the fluorescently labelled sample. The effective spatial resolution of LSFM can be improved with so-called multi-view imaging, where orthogonal views from two or more objectives are computationally merged, resulting in isotropic resolution (Preibisch et al., 2014; Swoger, Verveer, Greger, Huisken, & Stelzer, 2007).

To achieve the best image quality, the sample needs to be optically accessible from at least two directions, without significant differences of the refractive index between the imaging medium, the mounting medium, and the sample. To minimize refractive index changes, the sample is usually embedded in agarose and suspended in aqueous imaging medium. However, this embedding method restricts the range of manipulations with the sample: fast application of drugs or chemical stimulants to the sample becomes difficult due to the presence of agarose barrier and large volume of imaging medium. Furthermore, due to the labor-intense procedure of sample embedding and mounting, high-content screening becomes impractical.

Microfluidic chips are well-suited to tightly control experimental conditions, as the samples can occupy individual channels and be presented with inflow of nutrients and chemical stimulants, while metabolic waste is removed simultaneously. Furthermore, microfluidic chips allow conducting highly parallel experiments under nearly identical conditions. Conventional chips consist of a PDMS (polydimethylsiloxane) polymer layer, in which three-dimensional structures (e.g. channels) are formed and then enclosed with a glass coverslip.

The presence of glass coverslip is a major obstacle to the compatibility of LSFM with microfluidic chips. The orthogonality of objectives in a conventional light-sheet microscope requires that the glass coverslip is positioned at 45° angle to the objectives, which creates severe optical aberrations in both excitation and emission optical pathways, resulting in severe degradation of image quality. To address this problem, McGorthy et al (McGorty, Xie, & Huang, 2017) corrected aberrations in the detection arm using adaptive optics, but this approach did not generalize to multi-view imaging. Glaser et al (Glaser et al., 2019) used a solid-immersion lens (SIL) that minimized aberrations in both excitation and detection arms, also in single-view arrangement. Alternatively, several other SLFM designs (Dunsby, 2008; Sapoznik et al., 2020; Voleti et al., 2019; Yang et al., 2019, 2020) used one high-NA objective for both excitation and detection (single-view), at the cost of adding secondary and tertiary objectives used to re-image the tilted detection PSF onto the sensor.

In this work we present a daoSPIM, **d**ual-view **a**daptive **o**ptics **s**elective **p**lane **i**llumination **m**icroscope design which allows high quality imaging of life organisms without additional custom devices in front of the coverslip or the presence of secondary and tertiary re-imaging objectives. The optical aberrations caused by the tilted coverglass of the microfluidic device are efficiently compensated by deformable mirror. Moreover, due to the system’s symmetry, a single deformable mirror corrects aberrations in both optical arms simultaneously, greatly simplifying the use of microfluidic devices for LSFM imaging.

## Results

The light sheet is generated by scanning a focused laser beam through one objective, and collecting fluorescence through two objectives, while the sample is mechanically scanned through the excitation plane. The microscope is arranged in an open-top configuration (**Fig. 1a**) for maximal convenience, so that the coverslip-mounted sample is dipped into water-filled chamber from the top (**Fig. 1b**).

**Fig.1.**
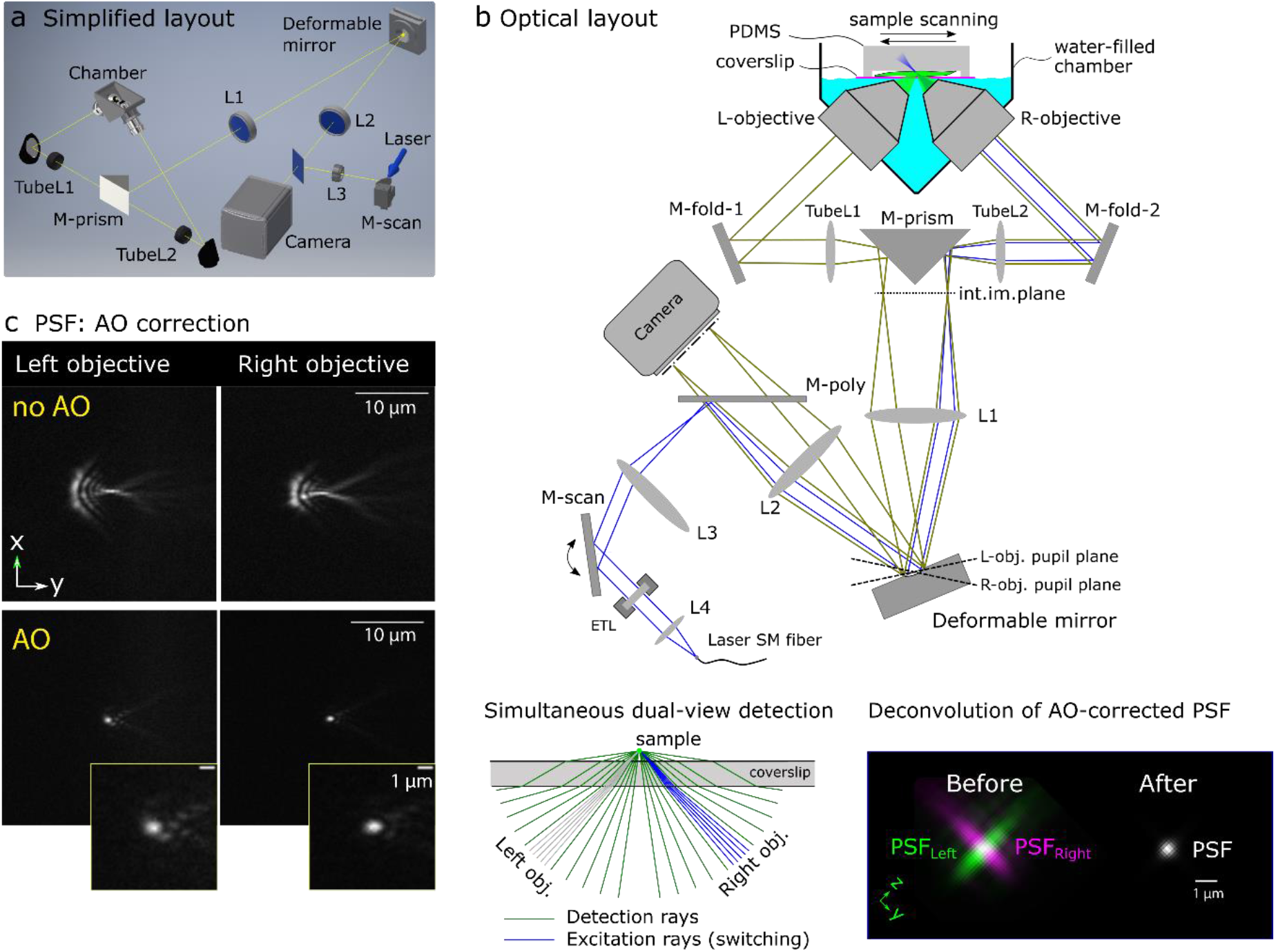
Microscope overview. (**a**) Simplified 3D model of the microscope layout. Scanning stage, mounts, and ETL module are omitted for clarity. (**b**) Optical layout with ray tracing: detection rays in green, excitation rays in blue. Abbreviations: *TubeL1,2*, tube lenses; *M-fold*, folding mirror; *M-prism*, prism mirror; *L1,2,3,4*, achromatic lenses, *M-poly*, polychroic mirror, *M-scan*, scanning galvo mirror, ETL, electro-tunable lens; *SM*, single-mode fiber from laser head. (**lower left**) Excitation can be directed into left (gray) or right (blue) objective, while detection is performed by both objectives simultaneously. (**lower right**) dual-view deconvolution of the AO-corrected PSF. (**c**) Fluorescent bead image in two views, before DM correction (no AO) and after correction (AO).

Due to the high cost of deformable mirror (DM), it is desirable to use a single DM for AO correction in both arms, which usually requires an optical switching mechanism that involves mechanical motion (e.g. flipping mirror). We avoided mechanical switching by adding a knife-edge prism mirror between the two arms (**Fig. 1b**, *M-prism*). We placed the deformable mirror at the intersection of the conjugated pupil planes of both objectives (**Fig. 1b** upper panel) to correct detection aberrations in both views simultaneously.

### Excitation path

The light-sheet is generated by scanning the excitation laser beam with the galvo mirror *M-scan* (**Fig. 1b** upper panel, blue rays). The laser beam starts in the joined path (laser out of *SM fiber -> L4 -> ETL -> M-scan -> L3 -> M-poly -> L2 -> DM -> L1*) and, depending on the galvo mirror voltage bias, reflects off the prism mirror M-prism into left or right arm. Thus, the galvo mirror *M-scan* serves two purposes: it switches the excitation between the arms, and scans the focused laser beam to create a light sheet plane in the active arm. An electro-tunable lens (ETL), placed before the galvo mirror, is used to adjust the light sheet axially.

### Detection path

The fluorescence light emitted by the sample, collected by both objectives simultaneously (**Fig. 1b**, lower left), is combined side-by-side by the M-prism. It then reflects off the DM, and is focused by lens L2 on the detection sensor (**Fig. 1b**, upper panel, green rays). The images from both arms are thus acquired side by side during the same exposure. Thereafter, dual-view deconvolution combines the two AO-corrected views into one, with improved spatial resolution (**Fig. 1b**, lower right). Due to symmetry of the optical system, the DM shape affects both views simultaneously and identically; thus aberrations in both views can be corrected by the same DM command (**Fig. 1c**).

### DM optimization algorithm

In order to find the best DM shape for aberration correction, we adapted the stochastic parallel gradient descent (SPGD) algorithm (Vorontsov, Carhart, & Ricklin, 1997), where we used dynamic gain control and an image-based metric that penalizes large-sized PSF in both arms simultaneously (Methods). We found that the algorithm robustly and rapidly converged to a minimal PSF in both arms (**Fig. 2**): cross-section FWHM(x,y) mean±std (0.44±0.01 μm, 0.75±0.01 μm, n=*10*) in right arm, and (0.49±0.05 μm, 0.74±0.02 μm, *n=10*) in left arm, measured with green fluorescent beads (diameter 0.17 μm, λ_em_ ~ 515 nm). The measured PSF in diffraction-limited conditions (beads in agarose sheet, water, no coverslip) was (0.44±0.003 μm, 0.49±0.04 μm, *n*=3). The theoretical diffraction-limited FWHM is 0.39 μm for NA=0.8 objectives used. The difference between the no-coverslip PSF and the theoretically expected size can be explained by the physical bead size and a noticeable astigmatism from the 4% agarose layer used for bead mounting. The fact that AO-corrected PSF is elongated in (y) is physically justified by the fact that a portion of NA in this direction is cropped by reflections off the coverslip (McGorty et al., 2017) and the steepest phase gradient occurs along this axis at the pupil edge (**Fig.2**).

**Fig. 2.**
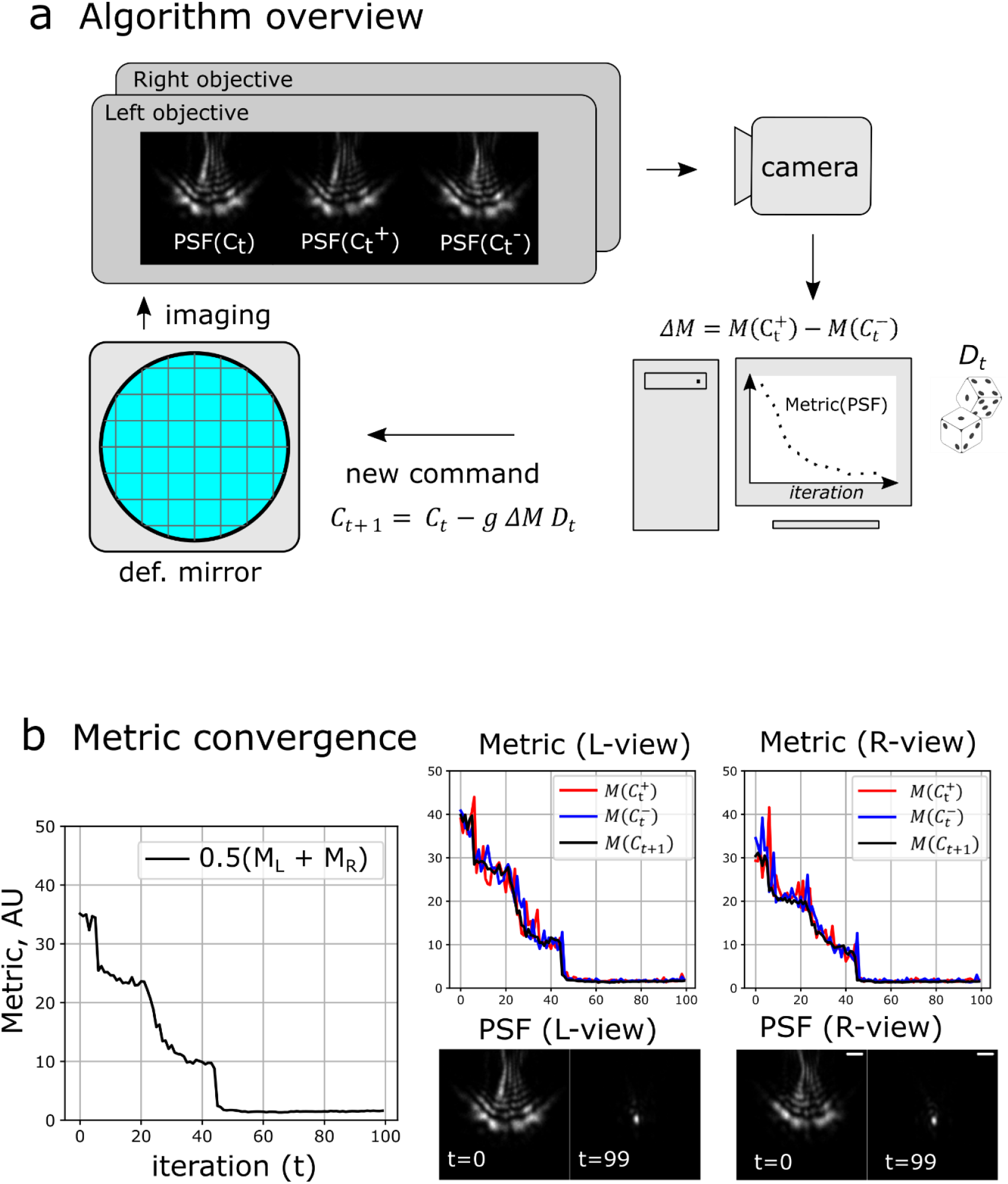
Algorithm for DM shape optimization. (a) Algorithm overview. At each iteration, three images are taken by the camera, and PSFs in each view are measured using metric *M*. (b) Total weighted metric convergence (left panel), and individual-view metric convergence (middle and right panels). Notation: *C_t_* is the DM command vector at time *t, g* is the gain, *D_t_* is the vector of random perturbations, *M*(*C_t_*) is the metric of PSF image obtained after command *C_t_* was applied to the DM. Scalebar 2 μm.

The DM optimization algorithm typically converged to a minimum in 50-60 iterations (2-3 minutes). The steep change of the metric (over 20x) was mitigated by the dynamic gain control, in which the gain increased inversely proportionally to the metric. This allowed having small gain at the beginning of optimization (stability) and high gain at the end (sensitivity). Once the optimal DM shape was found, it can be used for future experiments, provided that nominal coverslip thickness remained the same and the optomechanical alignment of the microscope was stable.

### AO-corrected detection FOV (green)

After finding the optimal (green-channel) PSF at the center of camera’s field of view (FOV), we measured detection PSF across the FOV to determine the spatial range (isoplanatic patch) across which a single DM command can correct aberrations. We found that the isoplanatic patch spans about 100 μm along Y axis (**Fig. 3**), in which green-channel PSF FWHM remains in the range (x) 0.46-0.47 μm, (y) 0.75-0.82 μm. With the camera sensor split between two objectives with system magnification 44.4x, each having 6.5 x 2048 / 44.4 / 2 = 150 μm wide maximal FOV, this makes useful detection FOV ca. 100 x 100 μm for each objective (with adjustment to FOV cropping at the prism mirror edge).

**Fig.3.**
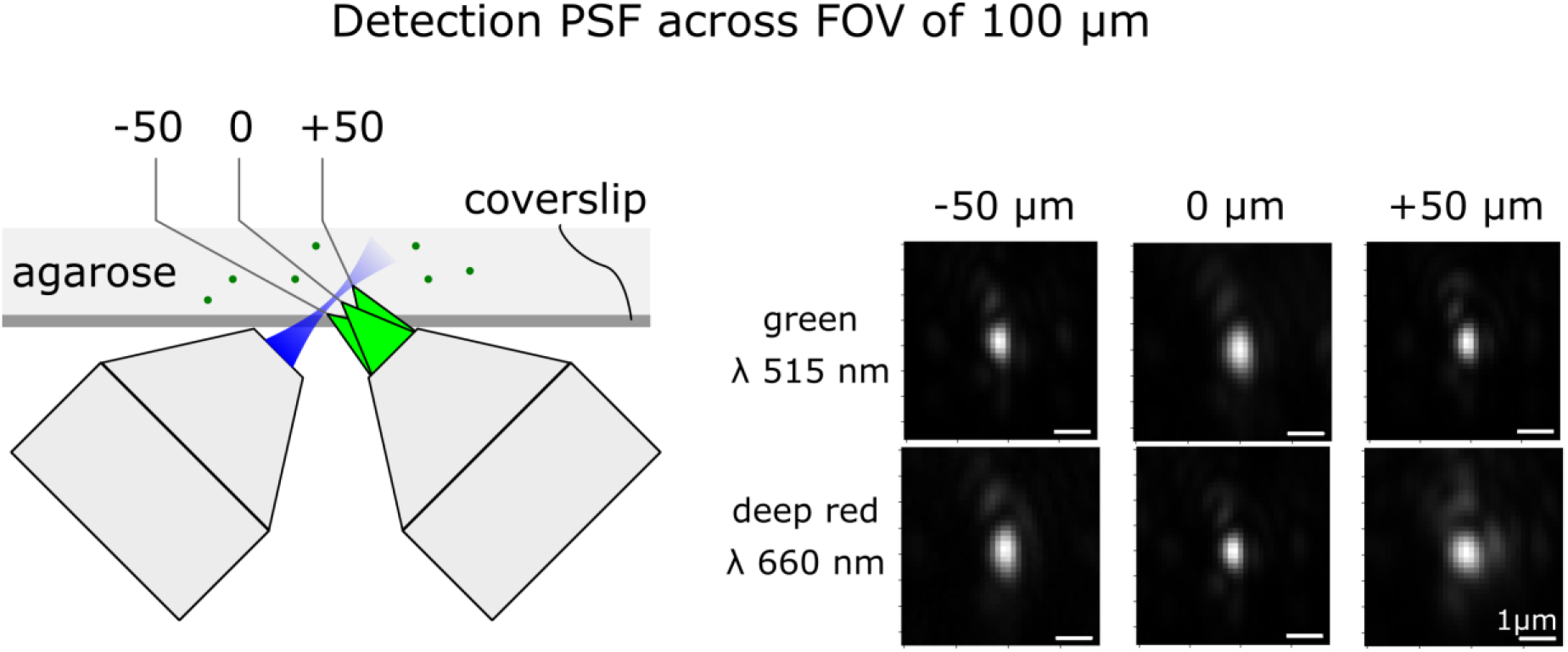
PSF measured using green and red fluorescent beads across field of view 100 μm. The DM shape was optimized using green beads. In green channel (ex/em 488/515 nm), FWHM(x,y) = 0.47, 0.75 (pos. 0); 0.46, 0.82 μm (pos. −50 μm), 0.47, 0.77 (pos +50 μm). In deep red channel (ex/em 638/660) FWHM(x,y) = 0.59, 1.03 (pos. 0); 0.59, 1.09 (pos. - 50 μm), 0.87, 1.01 (pos +50 μm).

### Multi-color detection

Because reflection changes the absolute optical path independently of the wavelength, the deformable-mirror based AO does not introduce additional chromatic aberrations beyond those already present due to refractive optical elements. Therefore, the DM shape optimized for green-channel PSF (λ_em_ ~ 515 nm) can be directly applied to image PSF in deep red channel (λ_em_ ~ 660 nm) (**Fig. 3**). The deep red PSF size scaled approximately proportionally to the emission wavelength (660/515 = 1.28x) across the 100 μm FOV: measured FWHM was (x) 0.59-0.87 μm; (y) 1.01-1.09 μm. Thus, the system was close to achromatic, and the DM shape optimized for one color channel can be directly applied to other channels.

### Dual-view PSF fusion

PSFs from the two AO-corrected views were computationally combined into a single PSF with a multi-view deconvolution algorithm using an additive update scheme (Methods). This reduced effective 3-dimensional PSF size FWHM(x,y,z) from (0.52, 0.79, 2.82) down to (0.26, 0.51, 0.62) μm, thus improving the resolution by factors (2.0, 1.5, 4.5) in (x,y,z) respectively (**Fig 4**). Note that x-axis is the one with smallest optical aberrations, y-axis has the highest aberrations, and z-axis in one PSF corresponds to y-axis of the other PSF after the two views are brought into the same coordinate system.

**Fig.4.**
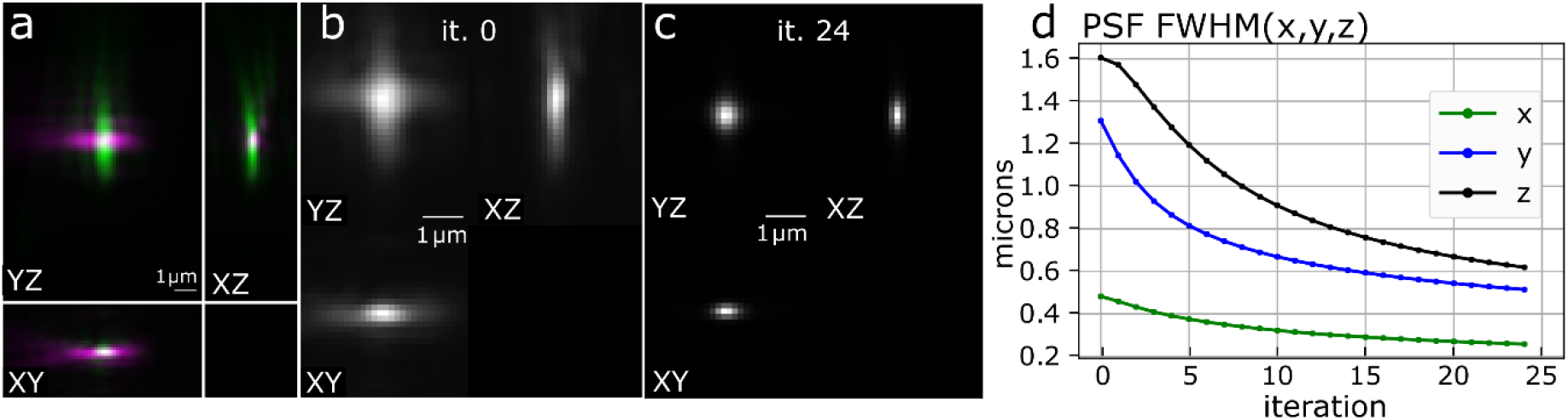
Dual-view PSF deconvolution, applied to AO-corrected PSF. (**a**) Overlaid AO-corrected PSF from the two views, acquired with ligh-sheet illumination. (**b**) Merged initial PSF, iteration 0. (**c**) Merged final PSF, iteration 24. (**d**) Changes of the merged PSF size (FWHM) during deconvolution.

### Light-sheet profiles

Because of the achromatic nature of deformable mirror correction, excitation light receives a phase shift identical in amplitude to detection light, but opposite in sign, and across a smaller pupil area (corresponding to light-sheet NA compared to detection NA). We thus expect that the detection-optimized DM shape also makes pre-shaping of excitation laser, so that after passing through the coverslip it becomes closer to diffraction-limited Gaussian beam. Indeed, we found that green-channel detection AO significantly improved the laser beam profile (**Fig. 5**). We visualized beam profiles for 488 nm and 561 nm excitation lasers as they illuminated green or red dye between the coverslips (fluorescein or rhodamine B, respectively). We found that AO-corrected beam waist was about 3.2 μm for both wavelengths, compared to coverslip-free 1.9 μm, and coverslip non-AO-corrected beam waist 5.2. Thus, green-channel detection AO correction provided satisfactory improvement in excitation beam profiles in both 488 and 561 nm excitation channels, albeit beam waist was thicker than in the no-coverslip case (**Table 1**).

**Fig.5.**
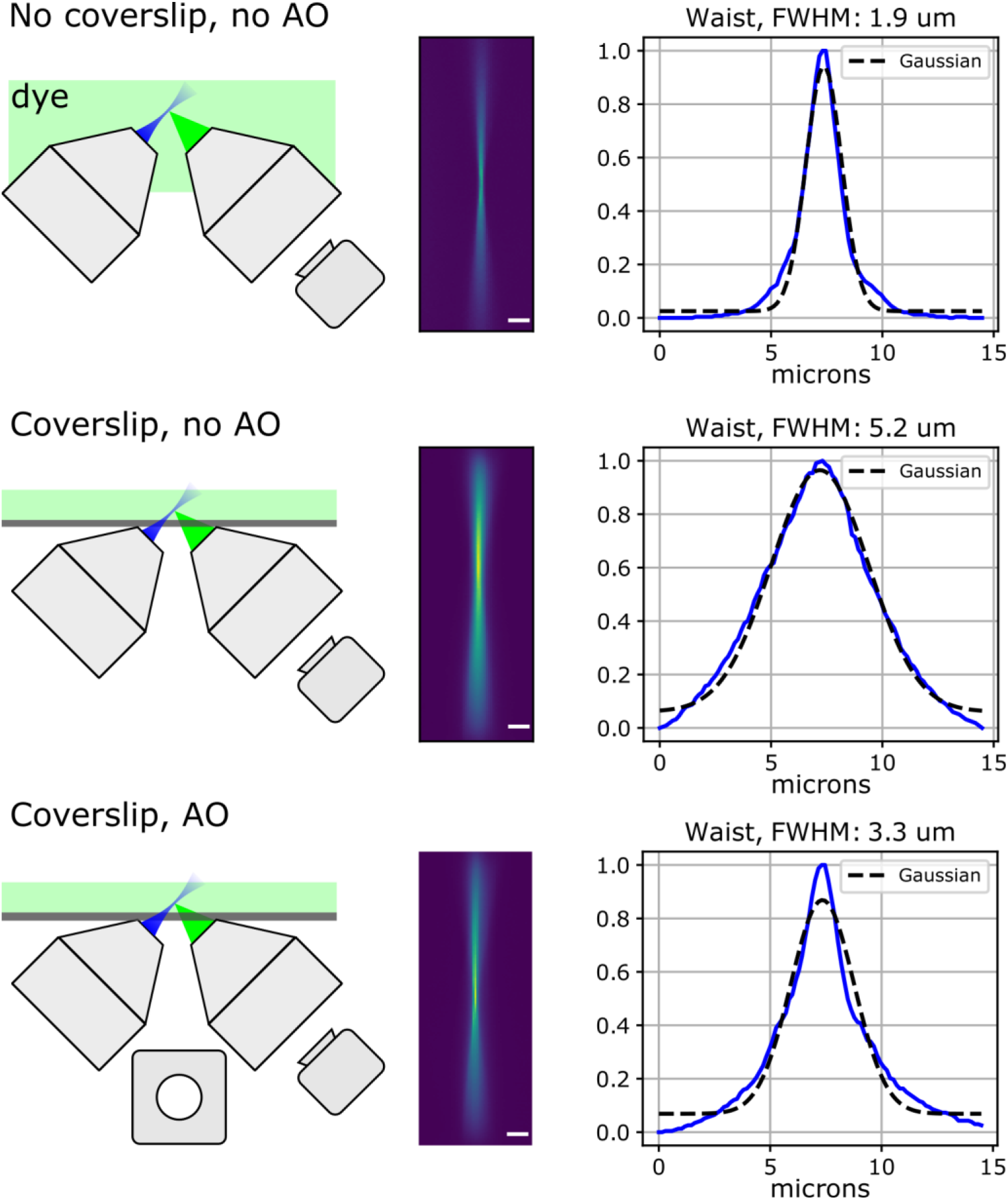
Light-sheet excitation beam properties. Image of the laser beam (NA 0.16) in fluorescein solution: no coverslip, with flat DM; after the beam passed through coverslip without AO (flat DM); with AO (DM shape optimized for detection PSF); Excitation 488 nm, emission ~515 nm. Scalebar 20 μm.

**Table 1.** Beam profiles in fluorescent dyes, with and without AO, compared to no-coverslip conditions.

| Beam parameter | Gaussian beam, dye bath, no coverslip, no AO |  | Coverslip and AO |  | Coverslip, no AO |  |
| --- | --- | --- | --- | --- | --- | --- |
| Excitation wl | 488 nm | 561 nm | 488 nm | 561 nm | 488 nm | 561 nm |
| Beam waist, $\mu\text{m}$ | 1.9 | 1.9 | 3.3 | 3.1 | 5.2 | 5.2 |
| Beam length (2 x Rayleigh range), $\mu\text{m}$ | 28.2 | 21.4 | 60.8 | 44.9 | 90.4 | 79.4 |

Measurement errors $\pm 0.1 \mu\text{m}$ (std).

### Imaging of *C.elegans* in microfluidic chip

We designed and manufactured microfluidic chips for imaging physically restrained *C.elegans* nematodes in tapered channels made of PDMS covalently bound to a high-precision #1.5H coverslip glass. The channels have a tip opening of 3 μm that prevents the nematodes from escaping. Nematodes with panneuronally expressing GCAMP6 (*AML32* line, dauer stage), were anesthetized and kept in the channels by applying mild static pressure.

After optimizing the DM shape using fluorescent beads, we imaged the head of an immobilized animal through the coverslip, along with individual fluorescent beads mounted in agarose to acquire PSFs for dual-view deconvolution (**Fig. 6**). Imaging with AO correction significantly improves image quality in each view over the image without AO, making individual neurons distinguishable. Computational merging of the left and right views by dual-view deconvolution further improved the image resolution, as expected from the nearly isotropic reconstructed PSF.

**Fig.6.**
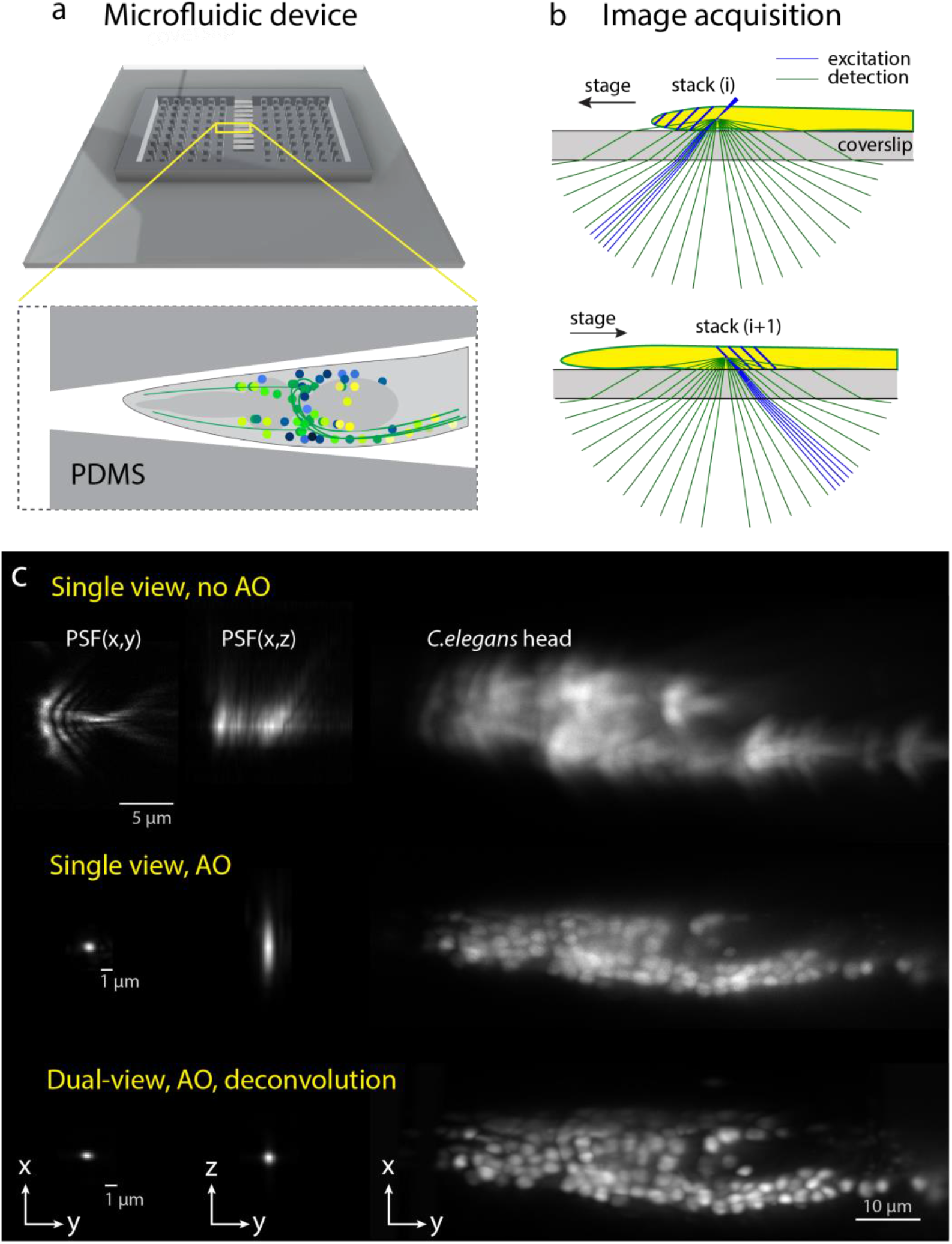
Dual-view imaging of *C.elegans* in microfluidic chip. (**a**) overview of microfluidic device on glass coverslip. (**b**) Stack acquisition scheme: right view is acquired when scanning the sample on one direction, left view when scanning in the opposite direction. (**c**) Point spread functions and single-view stack MIP of *C.elegans* head neurons acquired through the coverslip without AO (top row), with AO (middle row), and dual-view acquisition with AO, deconvolved and merged (bottom row).

## Methods

### Optomechanical design

The microscope has two water-dipping objectives (40X Nikon CFI APO NIR Objective, 0.80 NA, 3.5 mm WD) fitted in a custom 3D printed chamber (**Fig. 7a**) using custom-made silicone rubber O-rings. The objectives are fixed at 90° angle to each other on manual Z-axis translation mounts (Thorlabs SM1Z) to allow fine axial alignment. The microfluidic chip on a 22×22mm coverslip glass (#1.5H, VWR International, #630-2186) is held by small magnets in a sample holder and can be translated horizontally by a motorized stage (ASI FTP flat-top stage), and vertically by a manual translation stage (Newport M-461-X-M) with a small dove rail for quick sample holder release (**Fig. 7b**). The sample holder was custom-designed and manufactured from stainless steel (Protolabs). The 3D models of chamber, O-rings, and sample holder are available at https://github.com/nvladimus/daoSPIM along with other implementation details. The microscope components are listed in **Table 2**.

**Fig.7.**
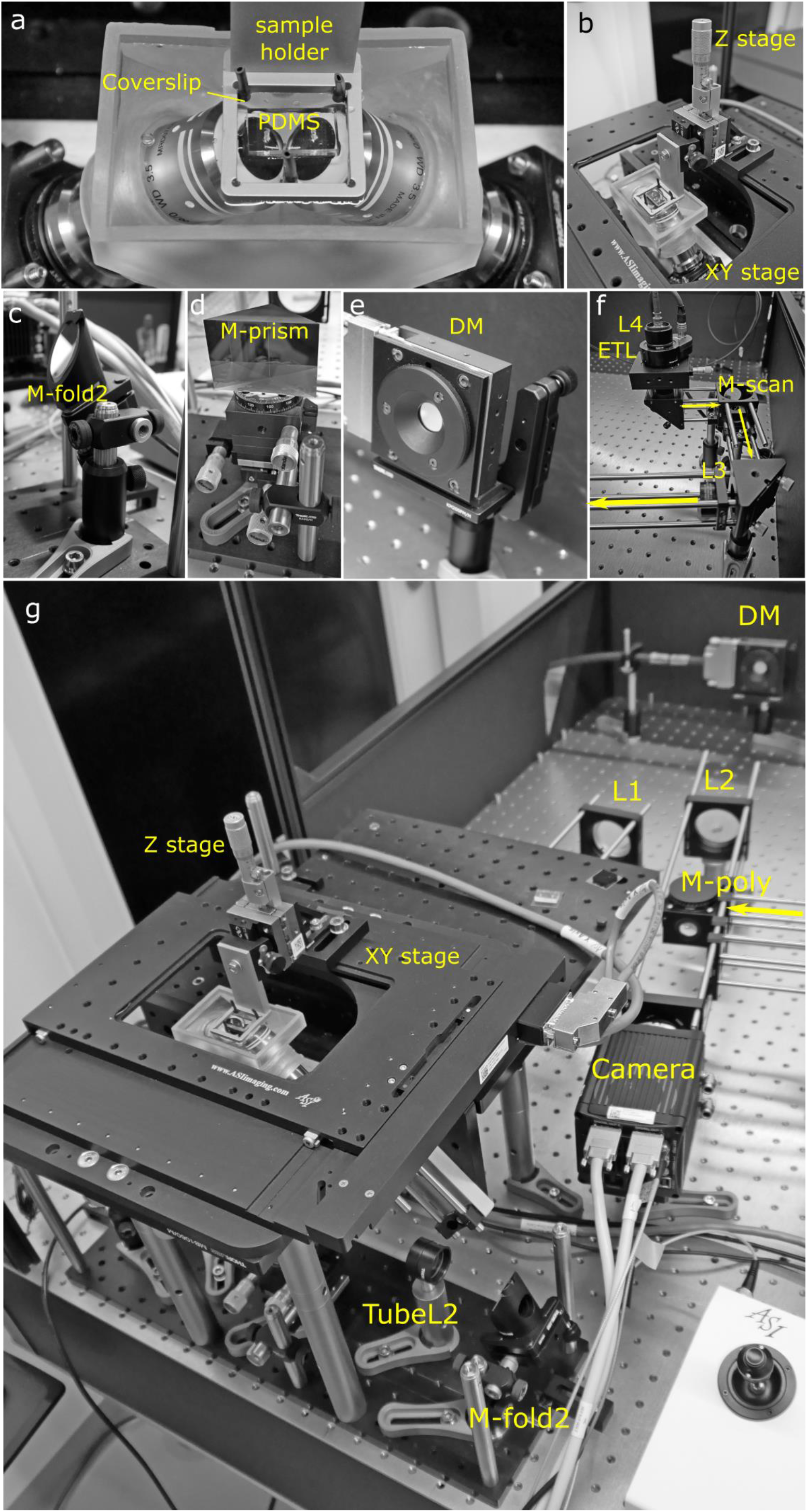
The optomechanical implementation. (**a**) The water-filled chamber with two objectives in open-top configuration. The microfluidic device is fixed inside the sample holder. (**b**) The sample holder is attached to motorized horizontal stage and manual vertical stage. (**c**) The folding mirror assembly. (**d**) The M-prism is mounted on the XYθ mini-stage. (**e**) The DM is fixed on a kinematic prism mount. (**f**) The laser scanning assembly, with yellow arrows showing laser propagation direction. (**g**) The system overview, with laser coming from the laser scanning assembly (yellow arrow).

**Table 2.**
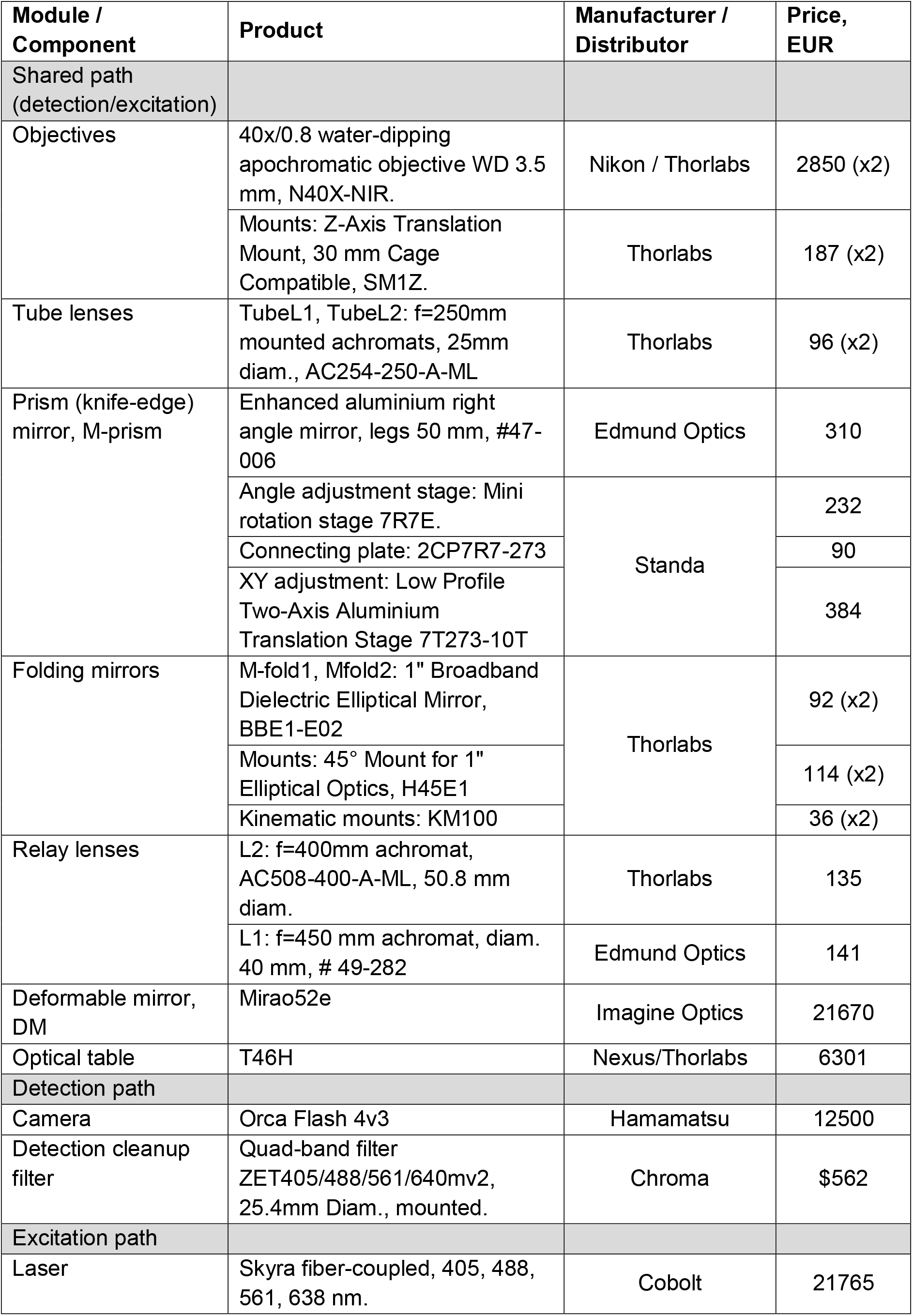
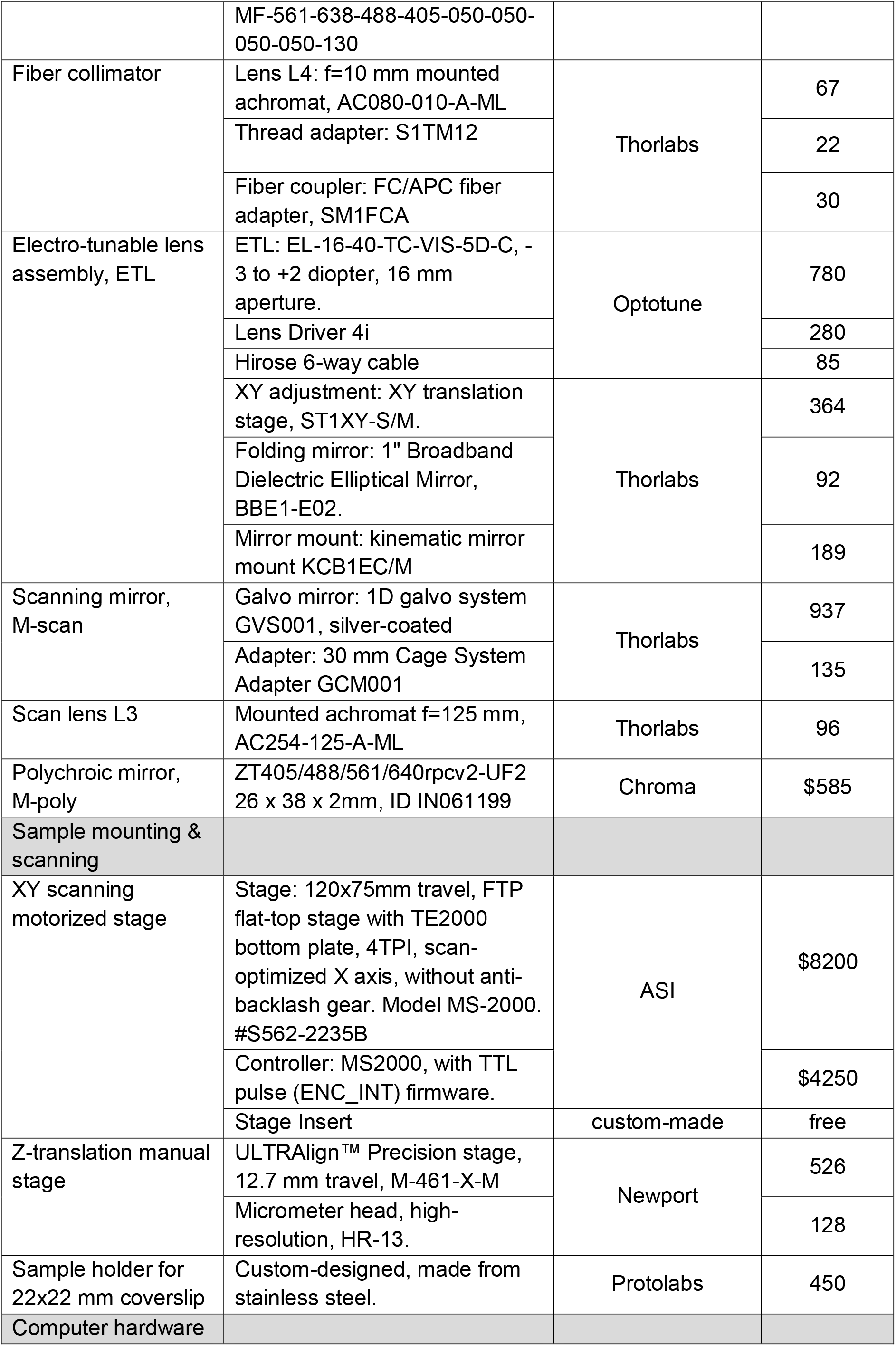
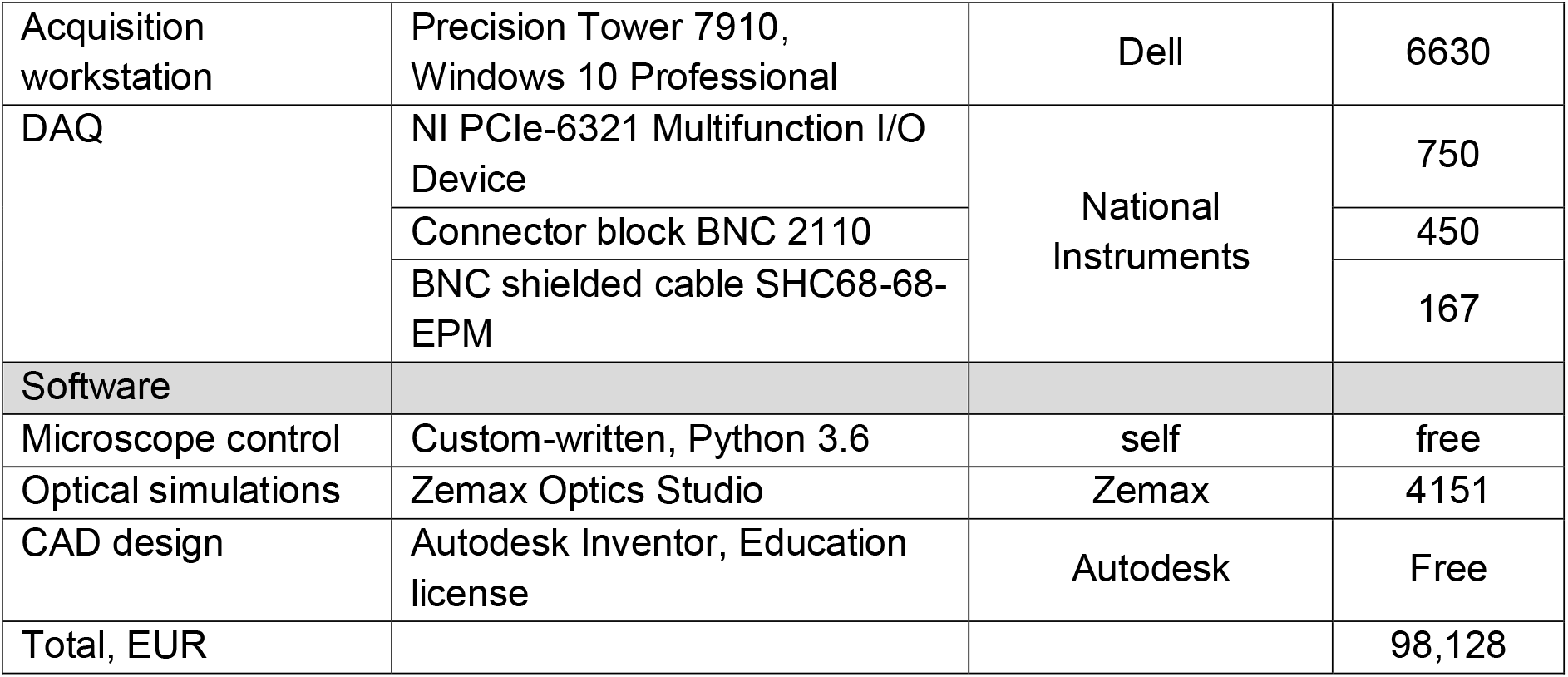
The microscope components.

The folding mirrors *M-fold1,2* (Thorlabs broadband dielectric elliptical BBE1-E02) are mounted on 45° mounts (Thorlabs H45E1) on top of kinematic mounts (Thorlabs KM100) and post assemblies for fine and crude adjustment, respectively (**Fig. 7c**).

The knife-edge mirror *M-prism* (Edmund Optics #47-006) was glued to an XYθ stage assembly (Standa 7R7E, 7T273-10T) (**Fig. 7d**). The DM (Imagine Optics, Mirao52e) was fixed (**Fig. 7e**) on a kinematic prism mount (Thorlabs KM200PM/M) and a custom adapter. The camera (Hamamatsu Orca Flash 4.3), polychroic mirror (Chroma ZT405/488/561/640rpcv2-UF2) and lens L2 (Thorlabs AC508-400-A-ML) were mounted in a 60-mm cage assembly.

The laser scanning module was mounted in a 30-mm cage assembly (**Fig. 7f**). The microscope overview is shown in **Fig. 7g**, with the laser scanning module omitted.

#### Deformable mirror

We used a continuous faceplate deformable mirror (Mirao52e, Imagine Optic), which has 52 electromagnetic actuators over an aperture of 15 mm. The back focal planes of both objectives were magnified with lenses *TubeL1/TubeL2* – and lens *L1* (**Fig. 1b**) while at the same time redirected toward the DM with *M-prism*. The DM was positioned at the intersection of (almost) overlapping conjugate pupil planes of both objectives.

### Iterative optimization

#### Metric

For DM shape optimization we adapted SPGD, **s**tochastic **p**arallel **g**radient **d**escent algorithm (Vorontsov et al., 1997) and minimized the following image metric

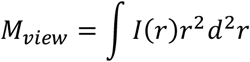

Here *I(r*) is the PSF image intensity (in left/right view) normalized to be within [0,1], *r* is the radius from PSF center. The SPGD should not be confused with SGD (stochastic gradient descent) commonly used in machine learning.

This metric choice was inspired by astronomy applications (Buffington, Crawford, Muller, Schwemin, & Smits, 1977): it penalizes large-sized PSF due to rapidly increasing *r^2^* term from the PSF center. The PSF center position was determined by thresholding the brightest pixels in the image (99% percentile) and computing their center of mass. After the main optimization, fine-tuning was sometimes necessary to further minimize FWHM(x,y) dimensions of the nearly-optimal PSF by another 15-20%. This was done by fitting PSF with 2D Gaussian profile and computing its FWHM(x,y).

All optimizations were performed on the metric averaged between the two arms,

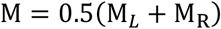

Since there are individual differences between the two optical arms, this metric balanced between them. However, the algorithm also performed well when one arm was given a preference (weights 0.75 : 0.25).

##### Initial conditions

The optimization had to start from relatively flat DM, so that initial PSF is not too distorted due to initial mirror shape. We initialized DM from factory-calibrated “flat” command or from all-zero command (non-flat DM shape), and found the latter giving more reproducible final optimized state, presumably because there is less initial bias compared to factory-provided flat command. Initial gain was set to 0.03, because values in the range of (0.01 to 0.05) gave the most stable convergence.

##### Iterations

At each iteration *t*, the current DM command C_t_=(c_1_, …, c_N_)_t_ (N=52 is the number of DM actuators) was perturbed twice in random directions by a perturbation command

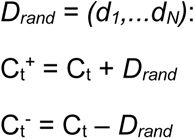

where components of *D_rand_* have randomly chosen signs and fixed amplitudes *abs*(d_i_) = 0.002 V (*i*=1..N). The perturbation amplitude of 0.002 V was empirically found to provide sufficient difference Δ*M* in metric between resulting images, but small enough to keep the algorithm stable. The new command was computed from the difference in cost function M between these two perturbations (Vorontsov et al., 1997):

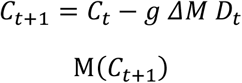

Here *g* is the gain parameter, 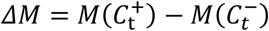 is the increment in cost function if commands C_t_^+^ and C_t_^-^ are applied to the DM. Gain *g* is a free parameter that critically affects the algorithm performance. Lower gain ensures stable convergence but slow speed of the algorithm, which is undesired due to increased sample exposure. High gain provides fast convergence but the algorithm can become unstable.

##### Dynamic gain control

In our case, the metric M changed considerably (by more than 20x) due to the highly distorted initial PSF image. To achieve both stable and fast convergence we made the gain adaptive (Finney, Persons, Henning, Hazen, & Whitley, 2014), so that it is scaled based on the initial and current metric values:

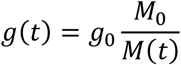

Here *g_0_* is the initial gain (0.03), *M*_0_ = *M*(*C*_0_) is the initial metric value with DM command *C*_0_ = (0,0,..0), and *M*(*t*) = *M*(*C_t_*) is the metric at iteration *t* with DM command *C_t_*. This dynamic gain adjustment ensured that the algorithm becomes more sensitive to very small changes of cost function M as the bead image approaches the diffraction-limited PSF size, which is only 3-5 pixels wide.

In all our experiments, the algorithm converged in 50-60 iterations (2-3 min). Small variations in final PSF shape can be attributed to the measurement noise and randomness inherent to the algorithm. Variations in the optimized DM command can also be attributed to the fact that the same PSF can be achieved by multiple commands due to invariance of PSF (x,y,z) translation that corresponds to DM tip, tilt, and defocus.

##### Regularization

In order to employ the left-right symmetry of the optical system and the expected left-right symmetry of the DM shape, we added a regularization step before updating DM command after each iteration, which averaged actuator commands with symmetrical positions relative to the DM middle:

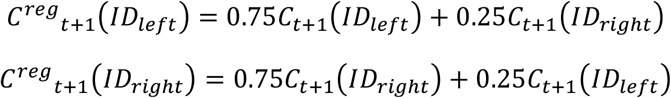

where *ID_left_* and *ID_right_* are the corresponding left and right actuator numbers (1..52).

We imaged green fluorescent beads mounted between two coverslips in 4% agarose and distilled water. The ROI in each arm was selected to contain a single bead (one bead in two orthogonal views), and was typically 100×100 px in size. The light-sheet laser was defocused to provide quasi-wide-field excitation, in order to avoid artefacts related to laser beam walking off the bead, since both detection and excitation path are controlled by the same deformable mirror.

A Python notebook with example optimization code is provided at https://github.com/nvladimus/daoSPIM.

### Optical simulations

To simulate the system wavefront at the pupil plane of objective accurately, one needs a model of the particular objective (in our case 40X Nikon CFI APO NIR Objective, 0.80 NA, 3.5 mm WD). Since the exact design is proprietary, we used three simplified models to assess the system wavefront (**Fig. 8**). All models were made in Zemax.

**Fig.8.**
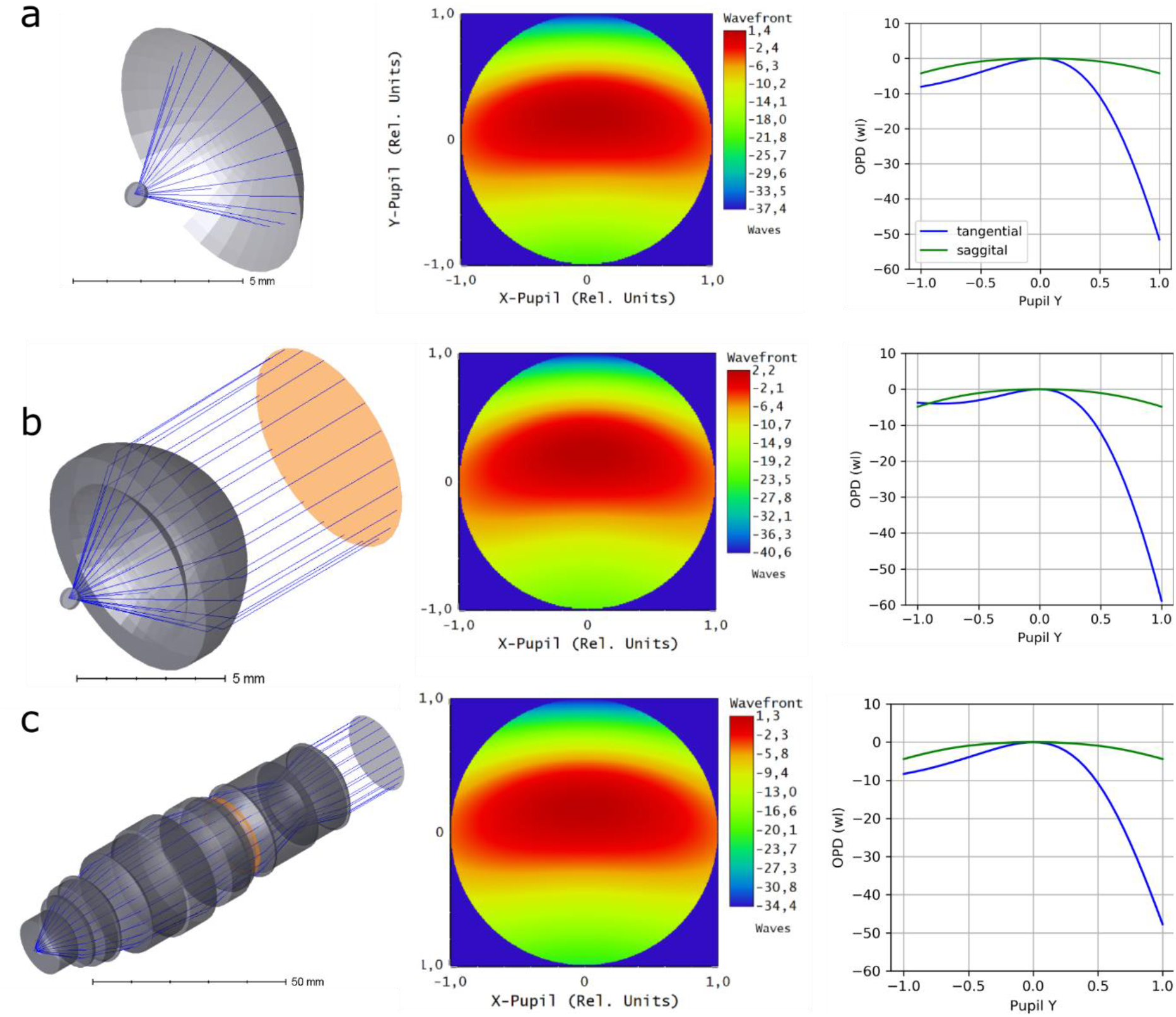
Ray tracing models and their predictions. (a) Model 1, coverslip tilted in water and spherical reference surface; (b) model 2, single aspheric lens in water; (c) model 3, Nikon 16x/0.8 water-dipping objective. Pupil planes are shown in orange. Second and thirds columns show simulated wavefronts relative to centroid ray (with y-tilt subtracted) and optical path difference to chief ray (with y-tilt preserved), respectively. OPD is shown in wavelength units (λ=520 nm).

#### Model 1: reference sphere in water

In the simplest approximation, we only inserted a tilted coverslip in water and plotted the relative phase map of rays at the reference spherical surface positioned 5 mm (focal distance of the objective) away from the point source (SI Fig 1). No other optical elements were used in this model, which minimized assumptions made. Coverslip thickness was 170 μm, tilt 45 degrees, glass type Schott D263-T-eco (n = 1.5233, Vd = 54.52).

#### Model 2: single aspheric lens objective

In this case, we modelled a 40x/0.8NA water-dipping objective by using a single lens of high-RI material (RI=2.41, similar to diamond), with spherical front surface (no refraction) and hyperbolic second surface, where all refraction occurs. This is a reasonable model of an objective for a monochromatic point source positioned in focus on the optical axis.

#### Model 3: using 16x/0.8NA objective model

We also used a detailed model of 16x/0.8NA Nikon water-dipping objective (kindly provided by D. Flickinger).

Despite the different assumptions of each model, their predicted aberrations agree between each other. The main aberrations are: vertical tilt, vertical astigmatism, defocus, vertical coma and vertical trefoil (listed in decreasing amplitude order), see **Table 3**.

**Table 3.** System wavefront RMS according to the models. All units are in wavelength (520 nm), the point source was on the optical axis.

| **Coefficient** | **Model 1**<br>**no lenses** | **Model 2**<br>**diamond lens** | **Model 3**<br>**Nikon 16x/0.8** |
| --- | --- | --- | --- |
| global y-tilt<br>subtracted | -14.2 | -18.3 | -13.3 |
| PV (to centroid) after<br>y-tilt subtraction | 38.8 | 42.8 | 35.7 |
| RMS after y-tilt<br>subtraction | 7.6 | 7.9 | 7.1 |
| Z3 (remaining y-tilt) | -7.1 | -9.1 | -6.7 |
| Z4 (defocus) | -4.8 | -5.1 | -4.7 |
| Z5 (o. astigmatism) | 0 | 0 | 0 |
| Z6 (v. astigmatism) | 5.3 | 5.4 | 5.0 |
| Z7 (v. coma) | -2.1 | -2.7 | -1.9 |
| Z8 (h. coma) | 0.0 | 0.0 | 0.0 |
| Z9 (v. trefoil) | 0.8 | 0.9 | 0.7 |
| Z10 (o. trefoil) | 0 | 0 | 0 |
| Z11 (spherical) | 0 | 0.04 | 0.06 |

#### Detection PSF measurement

We used the following fluorescent beads: 0.17 μm green (ex. 505, em. 515 nm) and deep red (630/660 nm) Invitrogen TetraSpeck™ Fluorescent Microspheres Sampler Kit; The beads were quickly mixed in 4% low-melting point agarose (Genaxxon Bioscience) at 70° C and sandwiched between two coverslips. The coverslips (#1.5H, VWR International, #630-2186) were sealed with dental glue (picodent twinsil^®^ speed).

#### Excitation laser beam profiling

For profiling 488 nm laser beam shape, we mixed fluorescein chloride (Sigma-Aldrich Chemie) with distilled water and sealed the solution between two coverslips with glue (picodent twinsil^®^ speed) and spacers of about 100 μm thick. For 561 nm laser beam profiling, we used Rhodamine B red (ChemCruz) in distilled water.

#### System synchronization

In order to capture optical sections of a sample with high speed and precision, one needs synchronization between illumination, camera, and stage motion with an accuracy of at least 0.1 ms with no jiter. Our system is driven by scanning XY stage (ASI FTP flat-top stage with TE2000 bottom plate, scan-optimized, 4 TPI) moving along the scanning direction in a serpentine mode, with TTL pulses fired every *ds* microns (0.5 to 2.8, depending on the sample) as the stage moves. The TTL pulses trigger the camera readout events (Hamamatsu Orca Flash 4.3 in SYNCREADOUT mode). The camera fires TTL pulses that indicate the global exposure period. The camera’s TTL pulses are captured by a digital counter task of an NI DAQmx board (PCIe-6321), which generates re-triggerable AO waveforms for galvo swipe and laser ON commands. The DAQmx task is created using PyDAQmx and runs continuously during the acquisition.

#### Laser switching between the arms

In order to switch the laser direction between the arms, one needs to add or subtract a voltage bias to/from the galvo-driving AO command after every *N*(images per stack) times. It was problematic to implement a re-triggerable AO task in PyDAQmx that meets this condition, so we added a custom PCB board (based on Teensy 2.0 MCU) that counts the camera exposure pulses and adds the left or right arm voltage bias to the galvo-driving signal in real time.

#### Control software

The control software was written in Python. In short, we wrote modular control modules for individual devices (camera, DM, light-sheet generator, scanning stage, ETL) which can be used as stand-alone programs (https://github.com/nvladimus/kekse). A top-level code (Python PyQt5) combined them into a specific workflow program (https://github.com/nvladimus/daoSPIM). The control software saved image stacks directly into HDF5 format using *npy2bdv* library (Vladimirov, 2020). The HDF5 format was chosen for compatibility with Fiji BigDataViewere/BigStitcher and Imaris software (Hörl et al., 2019; Schindelin et al., 2012).

#### Microfluidic device fabrication

The design consists of an inlet and outlet area, separated by 1.5mm long parallel, tapered channels to physically restrain nematodes. Standard microfabrication techniques were used to fabricate the microfluidic device (San-Miguel & Lu, 2013). Briefly, chip geometries were custom designed in CLEWIN software (WieWeb) and projected on SU-8 wafers using a MicroWriter ML3 (Durham Magneto Optics) optical lithography machine. Prior to photolithography, wafers were coated with SU-8 2010 (MicroChem) at a height of 15 μm (spin 10s - 500rpm, 100rpm/s acceleration; 30s - 1500 rpm, 30rpm/s). After exposing the wafer and baking according to SU-8 manufacturer’s instructions, wafers were developed by washing with PGMEA and isopropanol followed by a hardbake at 200°C for 20min. Master molds were used as negatives for polydimethylsiloxane (PDMS) replication. After applying PDMS (Sylgard 184, 1:10 ratio of curing agent and base), molds were baked at 110°C for 15 mins. PDMS devices were permanently bonded to cleaned #1.5 coverslips by a 30 second exposure to oxygen plasma. Bonded chips were baked on a hot plate at 75°C for one hour.

#### Worm handling

AML-32 nematodes expressing GCaMP6s in all neurons (Nguyen, Linder, Plummer, Shaevitz, & Leifer, 2017) were used for imaging. Dauer worms were extracted from starved plates that have been at 25°C for at least 5 days. Briefly, nematodes were washed off starved plates and washed twice with M9, before adding 1% SDS solution (dissolved in M9) and incubation for 30 mins at room temperature on a rotating wheel. Dauers were collected in M9 containing 1 mM tetramisol and loaded on microfluidic chips with the help of a 1ml syringe attached to a small piece of polytetrafluorethylen (PTFE) tubing.

#### Image acquisition

The sample was scanned at a constant speed through the light sheet generated by the scanning galvo mirror. This light-sheet generation mode (rather than via cylindrical lens) ensures that there is no motion blur, since the illumination is effectively stroboscopic: laser swipe time across the sample plane is 2 ms, the dwell time at any point is less than 0.1 ms, while the time interval between consecutive planes is 5-10 ms (limited by camera frame rate).

The image stacks were streamed into an HDF5 file along with an un-shearing affine transformation matrix that converted scanning stage intervals *ds* between successive images into corresponding *dz* intervals between planes 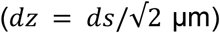. Thus, unsheared stacks could be visualized directly in BigStitcher without any post-processing.

#### Registration between left and right arms

Stacks taken by both objectives were resampled at the smallest available resolution (0.14625 μm/px) before further processing. Stacks taken by the right objective were transformed into the coordinate system of the left objective (transformations: inversion of X and Y axis signs, followed by rotation around OX axis by 90°). Once in the common coordinate system, both stacks were rotated +45° to fit a rectangular box with minimum background present. Registration pipeline included center-of-mass initial registration, intensity-based rigid registration, and ICP using points-of-interest (neurons).

#### Dual-view deconvolution

In order to compute fused image stack from two views, we used iterative dual-view Richardson-Lucy deconvolution (Lucy, 1974; Preibisch et al., 2014; Richardson, 1972; Wu et al., 2016) with experimentally measured PSF for each view.

For the registration and deconvolution we used BigStitcher (Hörl et al., 2019) and simpleITK (Lowekamp, Chen, Ibáñez, & Blezek, 2013; Yaniv, Lowekamp, Johnson, & Beare, 2018) software. Data analysis was performed using python *numpy* and *matplotlib* libraries (Hunter, 2007; Van Der Walt, Colbert, & Varoquaux, 2011).

## Discussion

Our AO microscope design demonstrates the possibility of dual-view light-sheet imaging through 45° tilted coverglass interface that separates objectives from the living sample. We designed our prototype system with the requirements of using off-the-shelf water-dipping objectives (Nikon 0.8/40x) and aqueous sample of extended lateral dimensions where the scanning is performed mechanically along a large spatial range.

In order to correct the extreme aberrations caused by the 45° tilted glass coverslip, we employ sensorless AO approach for PSF optimization with dynamic gain control. The latter speeds up convergence and ensures that the algorithm is not stuck in local minima. The algorithm currently optimizes PSF from a single fluorescent bead in the field of view. In future, optimization of extended sources (biological or synthetic fluorescent samples) would be preferred for optimization of detection PSF across the desired field of view. In principle, the use of AO is optional, since the aberrations due to the coverslip glass remain constant. Thus, the DM can be replaced by a fixed mirror with suitable surface curvature. However, DM has additional ability to compensate for errors in alignment and variations in the coverslip thickness and flatness.

Although a smaller tilt angle (e.g. 40°) generates less aberrations and allows for a better PSF correction (McGorty et al., 2017), we decided to use a 45° tilt in order to achieve symmetry between the two arms, necessary for multi-view imaging. We also dispensed with the use of pre-compensating cylindrical lens. Performing correction with a reflective element (DM) does not introduce chromatic aberrations and allows using it for multiple fluorophores and excitation lasers for simultaneous multi-color imaging. The measured thickness of excitation laser beam is further from the diffraction limit than the corresponding detection PSF, which requires further study.

The problem of open-top imaging through 45° tilted glass/polymer window was recently addressed by (Glaser et al., 2019) with a solid immersion lens, refractive index matching and multi-immersion objective design. Our method provides more aberrated PSF in each view and smaller field of view, but higher magnification, higher spatial resolution and it is compatible with live, dual-view imaging. The two methods solve the tilted glass problem in different application niches, and therefore complement each other.

A third approach to the problem of light-sheet imaging through glass is by using a single objective for both excitation and detection (Dunsby, 2008; Sapoznik et al., 2020; Voleti et al., 2019; Yang et al., 2019). More recent implementations achieve spatial resolution that matches high-NA spinning disk confocal imaging (Sapoznik et al., 2020), and multi-view imaging via beam inversion by two additional galvo mirrors (Yang et al., 2020). These systems require samples which have refractive index close to the imaging medium. The AO approach has additional flexibility to correct sample-induced aberrations, which can be useful in thick live samples that change over time.

The optical symmetry of our system provides several unique properties that are relevant outside of this particular application. Images from the two objectives are spatially separated on the camera chip, while their conjugate (Fourier) planes occupy the same spatial region, (where we place the DM), and their phases differ only by tilt. This allows simultaneous correction of aberrations, or generation of custom excitation beams, in two arms using a single AO element.

Finally, the low-cost beam splitting method we proposed (US Pat App. #16/679024) allows sharing camera, laser, and galvo scanner between two arms, thus significantly reducing the cost of existing diSPIM-like microscopes.

## Acknowledgements

NV was funded by EU Horizon 2020 MSCA Individual fellowship “Lightsheetelegans - In-toto imaging of C. elegans larval development using adaptive optics light-sheet microscopy” (grant no. 746497) and MDC Berlin; FP by Studienstiftung des Deutschen Volkes, BMBF grant no. 031L0142A and MDC Berlin; ZY by BCBB Support Services Contract HHSN316201300006W/HHSN27200002; AW and SP, MDC-Berlin, Additional support was provided by BMBF grant no. 031L0142A and, HFSP grant RGP0021/2018-102.

Authors thank D. Flickinger (HHMI Janelia JET) for sharing the Nikon water-dipping objective design, Dr. C.J. Bourgenot and Prof. J. Girkin (Durham University, UK) for help with code, and the Caenorhabditis Genetics Center at the University of Minnesota for providing *C. elegans* strains.

